# Ecology and conservation status of *Phoebe hainesiana* Brandis: a data deficit timber species from Indo-Myanmar biodiversity hotspot

**DOI:** 10.1101/2022.04.22.489227

**Authors:** Yengkhom Devajit, Lamabam Aashish, Benerjit Wairokpam, Vivek Vaishnav, Lokho Puni

## Abstract

*Phoebe hainesiana* Brandis (Lauraceae) is an economically significant timber yielding forest species endemic to North-East India in Indo-Myanmar Biodiversity Hotspot (IMBH). The tree exhibits significant commercial importance in timber industries due to its peculiar wood qualities. The timber’s strength, toughness, and durability make it in high demand for furniture making. Despite its significance, the botanical description of the species is not well documented due to the confined distribution of the species in IMBH region. The natural population is rare, but it is the ‘least concerned’ species in the IUCN Red List. Moreover, no taxonomical description of the species is available for its distinct identity among the other species from the genus. In the present study, we conducted a taxonomic, ecological, and forestry assessment of the species in its natural environment, researching the literature and conducting a field survey. A significant variation in intra-specific wood density (450 kg/m^3^ to 800 kg/m^3^) determines the scope of robust selection for the genetic improvement of the species. Its population has been severely fragmented due to over-harvesting in lack of sustainable management. According to our findings, the species is endangered due to a rapid decline in its population, a limited ‘extent of occurrence,’ and a small ‘area of occupancy.’ Among the other four species from the genus found in India, *P. hainesiana* can be identified based on its distinct lobes of the fruiting perianth. We recommend a sincere effort for the conservation of this key species of tropical evergreen forest in IMBH region.

## 1 INTRODUCTION

*Phoebe hainesiana* Brandis, synonym *P. goalparensis* (Lauraceae), also known as ‘Bonsum tree,’ is an endemic tree species to the Indo-Myanmar biodiversity hotspot (IMBH) region in North-East India (Brandis 1906). The timber obtained from the tree maintains toughness and durability and is lightweight, making it suitable for constructing household furniture, doors, windows, and other wooden stuff. It is denoted as ‘class-A1’ timber by the Indian government forest departments; hence the minimum sale price is second only to teak (*Tectona grandis* L. f.) wood. It is one of the most demanded timber species in the local furniture markets of North-East India. In its distributional range, the annual mean temperature varies from a minimum of 8°C to a maximum of 20°C with an annual rainfall of more than 2000 mm (Deb 1960). It prefers sandy-loam to clay soil types (Dowerah 1995) at a minimum altitude of 900 meters. It is a key species in montane sub-tropical climate forests of the IMBH region in India (Deb 1960). Due to high commercial demand, the species has been over-exploited through non-sustainable harvesting practices leading to fragmentation of the species population in the wild. Although the species is now a rare-sighting in- and outside of its natural habitat, it is a ‘least-concerned’ (LC) according to the IUCN - Red List. Despite regular utilization of the timbers obtained from the species by the indigenous people, the economic evaluation of the species has never been documented. The actual inventory about the utilization of its timber in regional and global trade is also not available. However, important information could be obtained because of the wide use of the species in the regional wood industries and trade in the Bhutan-India-Myanmar borders regions. Therefore, through this work, we documented all the available information of the species related to its taxonomy, population fitness, conservation status, and commercial significance reviewing available literature and performing field survey, sampling, and assessment.

## 2 METHODS

### 2.1 Distribution and taxonomical verification of the species

The information of the species distribution was obtained from the working plans of the state forest department, Government of India, literature available in the public domain (Google Scholar), GBIF (www.gbif.org), herbarium specimens of KEW (Herbarium, Royal Botanical Garden), and books published on Indian trees and timbers (Gamble, 1881; Brandis, 1906). This literature helped us also to understand the silviculture and management of the species. We conducted periodic field surveys throughout the natural range of the species in the IMBH region. We found its distribution in two natural patches *viz—*Senapati (latitude 25.424 N, longitude 94.290 E) and Ukhrul (latitude 25.043 N, longitude 94.235 E) forests of Manipur (Indian state within the IMBH region). The taxonomical identity and description of the species were verified through the Plant List (www.theplantlist.org) and herbarium specimen from KEW.

### 2.2 Ecological and conservation status of the species

We recorded the natural regeneration status of the species following the National working plan code (2014), applying a 0.1 ha size of quadrate. We recorded the distribution range, area of occupancy (AOO), extent of occurrence (EOC), population density, utilization pattern, and regeneration status of the species population following the Red List guidelines (IUCN, 2012) to assess the conservation status. We collected soil samples from 30cm depth to determine the soil type and measure the soil bulk density (weight/volume).

### 2.3 Assessment of the wood characteristics

We collected vertical and cross-sectional samples from the logs of the trees to study the physical properties of the wood. We also measured the basic wood density (dry weight/ volume) of the trees by extracting wood radial core samples from the breast height (1.37 m) and following the method described by Vaishnav (2018).

## 3 RESULTS AND DISCUSSION

### 3.1 Taxonomical and Botanical Description

There are more than 250 species recorded in the genus *Phoebe* Nees from the family Lauraceae distributed worldwide. Only 51 records have been accepted after taxonomical verification (The Plant List, 2013). Brandis (1906) documented seven species from the genus *Phoebe* in the Indian subcontinent. Two species *viz- P. lanceolata* (Nees) and *P. Paniculata* (Nees) were collected from central and peninsular India. Five species *viz- P. angustifolia* Meisn., *P. pallida* (Nees), *P. attenuate* (Nees), *P. tavoyana* Hook.f. and *P. hainesiana* Brandis were discovered in North-East India and some part of North India. Among these species from the Indian subcontinent, two *viz- P. lanceolata* and *P. pallida* have been found synonyms of the species *Ocotea lancifolia (Lauraceae)* and *Persea pallida* (Lauraceae) consequently. One species *P. augustifolia* Meisn. is yet to be verified. As per the latest available record, the taxonomical identities of the remaining three species than *P. haninesiana* have been accepted (The Plant List, 2013). The species *P. hainesiana* was firstly collected by H.H. Haines in 1893, and was named after Dietrich Brandis (Kew Herbarium Specimen No. K000778867 and K000778868). In 1916, a similar species named *P. goalparensis* was discovered by John Hutchinson (Kew Herbarium Specimen No. K000778865 and K000778866). Later, these specimens were authenticated as *P. hainesiana* Brandis (Kostermans, 1962). Although few published literature has cited the name of the species *P. hainesiana* Brandis as *P. goalparensis*, it is a synonym of *P. hainesiana* Brandis itself.

The plant species of the Lauraceae are known for their complicated taxonomy and are challenging in identification morphologically (Yang et al., 2021). Species from the genus *Phoebe* Nees are also complicated to identify well based on floral traits alone. Shu et al. (2008) provided a taxonomy description of the Phoebe Nees species and taxonomic keys for their identification. Brandis (1906) documented the taxonomical description of the Indian species from *Phoebe* Nees (Table 1). We found that *P. hainesiana*Brandis can be distinctly identified among other species of *Phoebe* in India, only based on the six tepals (three inners and three outers) of perianth attaching the fruit to peduncle (Figure 1). In *P. hainesiana*, lobes of the fruiting perianth of the species are coriaceous and are not appressed.

**TABLE 1.** A comparative taxonomical key description of Indian species from genus *Phoebe* Nees (Brandis 1906)

| Descriptions | *Phoebe* species in India |  |  |  |
| --- | --- | --- | --- | --- |
|  | A. Lobes of fruiting perianth appressed, rigid, almost horny |  |  | B. Lobes of fruiting perianth coriaceous, not appressed. |
|  | *P. paniculata* | *P. attenuata* | *P. tavoyana* | *P. hainesiana* |
| **Habit** | Moderate size tree | Large tree | - | Large tree |
| **Stem/shoot character** | Young shoot rusty *tomentose | Branchlets *stout, and the shoot is rusty tomentose with long soft hair | - | The stem bark is thick and dark in color |
| **Leaf** | *Pubescent beneath and midrib of above; elliptical or obovate elliptic; blade narrowed into petiole | Petiole, the underside of the leaf is rusty tomentose with long soft hair; oblanceolate; blade narrowed into a petiole; tertiary vein parallel | Membranous leaf, *acuminate with often scattered minute hair beneath; blade narrowed into a petiole; secondary vein arching with shorter intermediate vein or inter-secondary vein | *Glabrous, pale beneath and clustered at the end of a branch; oblanceolate to obovate; secondary vein parallel, prominent beneath |
| **Flower/ Inflorescence** | Inflorescence panicle and flower pubescent on the long slender peduncle, partly enclosed by persistent perianth | Inflorescence rusty tomentose with long soft hair, spreading panicle; peduncle stout | Inflorescence panicle and slender | The inflorescence is panicle; peduncle stout; perianth grey, softly tomentose on both side |
| **Fruit** | - | Narrowly ellipsoidal; *fruiting calyx | - | Black in color, fleshy, and base enclosed by *campanulate perianths |

\*Tomentose- Plant hair flattened and matted, Pubescent- Covered with soft short hair, Glabrous- Not hairy, Stout- Large, bulky, Acuminate- Tapering to a point, Fruiting calyx- Calyx extended to support fruiting, Campanulate perianth- Widened perianth.

**FIGURE 1.**
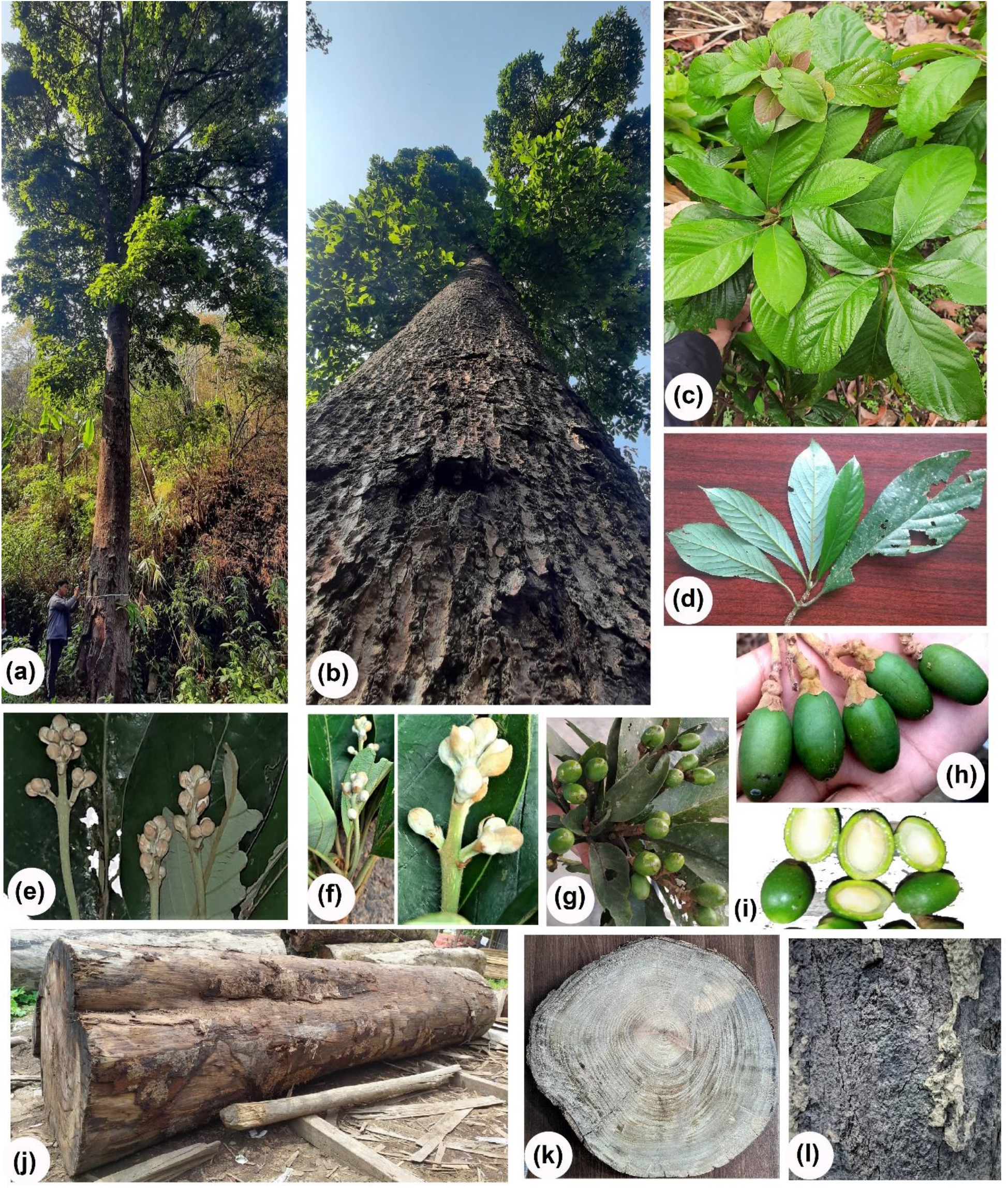
Plant parts of *P. hainesiana* (a) A moderately aged tree, sampled for the study, (b) Branching and clear bole, (c) Phyllotaxy, (d) dorsiventral leaves, (e) and (f) flowers, (g) and (h) fruits, (i) seed section, (j) log of felled tree, (k) cross-section of the log, (l) bark surface.

### 3.2 Phenology and Floral Description

*P. hainesiana* Brandis is an evergreen tree that flowers during April -June, and fruits during and July-November. Leaves are lanceolate, obovate, or ovate, rarely obtuse at the base, and are glabrous, 9-12 lateral veins on both halves of the leaf (Dowerah, 1995). Venation of leaf is pinnate camptodromous with secondary vein having brochidodromous festooned. The secondary vein divergence angle is acute and narrow. Furthermore, tertiary veins are arranged in a randomly reticulate manner. Tracheoids are present in the leaf (Chodankar & Vaidya, 2021). Flowers are bisexual, perigynous, and the ovary is covered with a cup-like structure at the base (Kundu, 2012; Pal, 1979). The panicle inflorescence is loose (lax panicle) in a long peduncle, pedicel, or stalk is about 0.5cm long, minutely pubescent, and presents minute bracteoles. Flower buds are ovoid and acute. Perianth segments are ovate, obtuse, puberulous outside, appressed, and villious/shaggy inside towards the base. Stamen is slender, and anther is four celled. The ovary is depressed, pubescent, and spherical/globose (Dowerah, 1995; Kundu, 2012). Fruits are berries, aromatic, ellipsoidal in shape, glabrous, blackish when ripening, and mesocarp is thin (Bisht et al., 1999; Kundu, 2012). The Young branches of the tree are minutely pubescent (Kundu, 2012). The bark is dark or blackish-grey in color (Brandis, 1906), and the surface is reticulately furrowed.

### 3.3 Ecology

*P. hainesiana* Brandis is a tall evergreen tree, which can attain a height of 30 - 35 meters. In the Indo-Myanmar border state Manipur (India), local peoples call it ‘Uningthou,’ which means ‘king of trees’ because of its giant size in the forests. We found it distributed in the altitudinal range of 1400 m to 2100 m, temperature range of 4 °C (in the coldest month of the year) to 30 °C (in the hottest month of the year), and annual precipitation of more than 1500 mm. It can grow on soil ranging from sandy loam to clay. It is a shade bearer species, and young plants cannot withstand direct exposure to sunlight (Dowerah, 1995). However, at the sapling stage, it requires light for fast growth. The natural populations of the species are distributed in the tropical and sub-tropical evergreen forests of the IMBH region confined to Indian territory. In Manipur, the natural patches of the species were found in montane sub-tropical climates in the plant climax community of *Saurauia – Phoebe – Beilschmiedia* (Kangpokpi hills) and *Laurus – Melia – Bauhinia* (Koubru hills) units (Deb 1960).

### 3.4 Reproductive Behavior and Artificial Regeneration

We observed good natural regeneration of the species through fallen seeds under tree canopy only in one patch. Coppicing was also observed in an aged tree of *P. hainesiana*. The species perform well in natural and artificial regeneration only if the seeds get the optimum conditions after falling. In November-December, seeds are dispersed by birds and animals, who eat the fruit-pulp and leave the seed. The seed is recalcitrant (Bisht et al., 1999; Kundu, 2012) and requires moisture and shade for regeneration within a few weeks of its falling. It exhibits hypogeal germination. The seeds do not remain viable for more than two-three months.

For storage, the recalcitrant seeds of the species cannot withstand desiccation. The seed remains viable for six months at normal conditions (temperature range of 15°C - 30°C). Under controlled conditions with 32 % to 34% moisture content and 5°C to 15°C temperatures, the seed can be stored for two years. Germination of the seed is adversely affected if the moisture content goes down below 29% (Kundu, 2012). A 70% germination has been observed in the seeds stored for three months (Bisht et al., 1999). The seeds can be treated with 0.2% Bavistin before storage to avoid fungal infection (Kundu, 2012).

For artificial regeneration, seeds are given pre-sowing treatments of de-pulping, scratching, and soaking overnight in 0.05% gibberellic acid (Jana & Singh, 2017). Mechanical de-pulping followed by drying for 8 hours or mechanical de-pulping followed by soaking in normal water for 24 hours also give an excellent germination percentage (Jana & Singh, 2017). The hypogeal germination of the seed typically takes25 days to 90 days or more (Bisht et al., 1999; Kundu, 2012). For propagation, raised nursery-beds are prepared, and seeds are planted at a depth of 1.5 cm to 2.0 cm (Dowerah, 1995; Kundu, 2012), preferably in November after drying (Bisht et al., 1999). Seedlings cannot withstand direct exposure to sunlight (Dowerah, 1995). Therefore, nursery beds are preferred in the shade.

After four months of sowing, at the stage of three leaves, seedlings can be transplanted. After six months of transplantation, the seedlings can be established in the field. For this species, stump planting is found more reliable to some extent than direct sowing (Bisht et al., 1999; Kundu, 2012). Proper care should be taken up to the establishment stage for a good survival percentage of the seedlings (Bhatt et al., 2010; Kundu, 2012). Root conditioning treatment is an effective treatment for better performance of the seedlings. Undercutting, base cutting, and wrenching treatment prevent root coiling and positively impact the seedlings’ survival rate, height, and diameter. At the same time, out-planting wrenching encourages the lateral root system, which helps in nutrient absorption, suitable for seedling establishment (Jana et al., 2017).

### 3.5 Silviculture and Silvicultural System

*P. hainesiana* is planted through diffuse and tunnel planting techniques in natural forests (Dowerah 1995). In diffuse planting, the seedlings are transplanted, maintaining a distance of 3 to 4.5m or more, in the natural forest under a moderately open canopy. The tunnel planting is generally adopted for the restocking of the forest. In this method, tunnel-like strips are made at an interval of 3 to 4.5m by removing shrubs undergrowth and some middle canopy trees. Seedlings are planted out along the tunnel 3 to 4.5m apart. Eventually, the overhead canopy opens up gradually with the progress of seedlings (Dowerah 1995). In natural forests of lower Eastern-Himalaya, the species is managed under shelterwood or selection system mainly for timber production (State of Forest Genetic Resource in India: A country report, 2012).

### 3.6 Uses and Utilization

*P. hainesiana* is mainly utilized for its excellent quality timber in wood industries. Due to its peculiar wood grain and structure, it is also called ‘Assamese teak’. The wood log of the species is utilized for making furniture, doors, almirah, planks. (Dowerah, 1995). We found that the species’ round log costs a minimum of USD 12 / cubic feet square, and the plank costs a minimum of USD 15 / cubic feet square in the Indo-Myanmar border region. The species has sacred significance among the ethnic communities of Manipur. The wood log from the species was used to make boats for the royal family of the princely state of Manipur in India. The tree’s bark contains antioxidants such as flavonoids and polyphenols, which are crucial in maintaining standard health requirements, and steroids, tannin, and saponin (Narah et al., 2016). Nevertheless, the species is majorly utilized in timber industries only.

### 3.7 Wood Properties

*P. hainesiana* Brandis exhibits diffuse-porous wood with indistinct growth rings. Vessels are solitary and distributed radially in multiples (2-7) and clustered (Singh et al., 2015). The species’ wood is considered under ‘class-II’ for its durability, which means this wood has a life span of five to 10 years (Sundararaj et al., 2015), preventing decaying pathogens. However, decay of dry wood with mottled sponge rot (caused by *Ganoderma applanatum* Pers.), white fibrous rot (produced by *Tramates serpens* Fr.), white spongy rot (caused by *Trametes corrugate* (Pers.) Bres.), and brownish pocket rot (caused by *Polyporus gilvus* (Schwein) has been documented (Dowerah 1995; Kundu, 2012).

The wood from the species seasons well, within a short period, without seasoning defects. It has moderately refractory properties. The basic wood density of the wood varies between 450 kg/m^3^ to 800 kg/m^3^. We found a significant difference in intraspecific wood density. It confirms the need for robust selection for genetic improvement of the species.

## 4 Threat to the species population

The species’ population has been over-exploited due to a lack of proper harvesting techniques and a standard rotation age. Due to significant commercial demand, the trees are illegally cut and utilized in local furniture industries. Shifting cultivation (jhum cultivation) or ‘slash and burn’ practiced by indigenous communities is also a threat to the species population. Along with this, the absence of natural regeneration through seeds in the natural populations of the species and lack of artificial regeneration protocol also led to population fragmentation, declining genetic variability, and intra-specific gene flow. The ecological niche of the species population is confined to hill areas with specific soil types; therefore, the fitness of the population is also at threat of projected climate change.

## 5 Conservation Status

*P. hainesiana*Brandis (synonym *P. goalparensis*) had been enlisted under the least concerned (LC) category of the IUCN Red List. During our assessment, we observed a population reduction of more than 70% in the last ten years and more than 90% over the length of three generations. The EOC is limited to less than 5000 km^2^ with an AOO of less than 10 km^2^ in the niche of tropical evergreen woods and a specified altitudinal range. The species population is heavily fragmented, containing less than 100 mature individuals in any of its sub-populations. The species’ geographical distribution is limited to a restricted population with less than 50 mature individuals per hectare of area. With available assessed data, the conservation status of the species may be considered under the ‘endangered’ category by the IUCN Red List.

## 6 Conclusion

IMBH region covers one of the least explored forests. *P. hainesiana* Brandis is a prioritized species of the region for economic utility and conservation value in India. The state forest department, Arunachal Pradesh (Government of India), has established a seed orchard for genetic improvement of the species. *P. hainesiana* is among the 153 species shortlisted and prioritized by APFORGEN (Asia-Pacific Forest Genetic Resources Programme), FAO (Food and Agriculture Organization), FGRMN (Forest Genetic Resources Management Network), and ICFRE (Indian Council of Forestry Research and Education) for genetic improvement and conservation (State of Forest Genetic Resource in India: A country report, 2012). This endemic species yields valuable timber, which is highly utilized in regional timber industries. The locals have overexploited it without sustainable harvesting practice, which has caused a significant reduction in its occurrence. There is an urgent need to select the superior genetic resource of the species and develop a rapid propagation technique for its mass multiplication to fulfill the need of the timber industries for the long term. To the best of our knowledge, the present work is the first account of the species’ taxonomic description, ecology, silviculture, and conservation status. Nevertheless, the taxonomical ambiguity among the species from the genus can be resolved through the DNA marker-based phylogenetic assessment. For ecological restoration of the species, there is a need to develop its propagation methods through biotechnological interventions. It is also recommended that the conservation status of the species should be in the ‘endangered’ category in the IUCN Red List.

## ACKNOWLEDGEMENTS

The authors are grateful to the State Forest Development Agency and State Forest Department, Government of Manipur, India, for fund support and permitting access to the forests of Manipur. The institutional support from the competent authority, Manipur University, is also gratefully acknowledged.

## CONFLICT OF INTEREST

The authors declare no conflict of interest among them.

## AUTHOR CONTRIBUTION

All authors contributed equally to the literature survey. V.V. is the research project’s principal investigator, who designed the study and updated the manuscript before final submission. Y. D., L. A., and B. W. carried out the field sampling, survey, and laboratory work. Y. D. prepared the first draft of the manuscript.

## Notes

### Competing Interest Statement

The authors have declared no competing interest.

